# Hepatitis E ORF2 blocks trophoblast autophagy to induce miscarriage via LC3B binding rather than PI3K/Akt/mTOR pathway suppression

**DOI:** 10.1101/2025.09.19.677307

**Authors:** Yinzhu Chen, Yifei Yang, Qianyu Bai, Xinyuan Tian, Chaoyu Zhou, Xuancheng Lu, Tianlong Liu

## Abstract

Hepatitis E virus (HEV) is a zoonotic pathogen that can infect pregnant women and cause adverse pregnancy outcomes, including miscarriage and preterm delivery. The previous study demonstrated that HEV genotype 3 (HEV-3) inhibits complete autophagic flux in both mouse placental tissue and human trophoblast cells (JEG-3), evidenced by reduced expression of ATG proteins (including LC3, Beclin1, ATG4B, ATG5, and ATG9A) and accumulation of p62. However, the specific regulatory pathway involved remains unclear. Thus, eukaryotic expression vectors for HEV open reading frames (ORFs) were constructed, and ORF2 and ORF3 proteins were transiently overexpressed in JEG-3 cells via liposome transfection.. While both ORF2 and ORF3 significantly reduced LC3B protein levels (*p* < 0.01), only ORF2 induced p62 accumulation (*p* < 0.01), indicative of autophagic inhibition, which indicates that ORF2 was the key viral protein mediating autophagy suppression in JEG-3. The results of WB and RT-qPCR show that ORF2 suppressed the PI3K/Akt/mTOR pathway while enhancing nuclear translocation of TFEB (*p* < 0.01) and AMPK phosphorylation (*p* < 0.01), suggesting paradoxical activation of upstream autophagy regulators. Through co-transfection of mCherry-LC3 with ORF2, co-localization studies, and AlphaFold 3-based intermolecular interaction predictions, we proposed that ORF2 directly binds LC3B to block autophagosome formation. Finally, co-immunoprecipitation confirmed physical interaction between HEV ORF2 and LC3B, elucidating the molecular mechanism of HEV-induced autophagy suppression in trophoblasts. These findings reveal the molecular mechanism by which HEV inhibits autophagy leading to miscarriage in mice, providing new insights into HEV-induced reproductive damage.

## Introduction

Autophagy serves as a cytoprotective mechanism that is activated in response to intracellular stressors such as protein aggregates, damaged organelles, or pathogenic invaders, thereby preserving cellular homeostasis and functionality[1]. The overall process of autophagy is referred to as the autophagic flux, which includes three stages: autophagy initiation, autophagosome formation and maturation, and autophagosome-lysosome fusion and degradation, coordinated by autophagy-related (ATG) proteins and involves the mobilization of various cellular components[2]. The PI3K/Akt/mTOR signaling pathway serves as the core upstream regulatory pathway of autophagy, negatively modulating the metabolic processes of synthesis and degradation of autophagy-related genes[3]. The mTOR pathway serves as a critical regulator in placental trophoblasts[4], with mTORC1 being the predominant pharmacological target for modulating placental autophagy[5]. As the master transcriptional regulator of autophagy and lysosomal biogenesis[6], TFEB is dynamically controlled in trophoblasts through mTORC1-dependent phosphorylation[7,8]. Additionally, AMPK is an evolutionarily conserved serine/threonine protein kinase that initiates downstream autophagy by phosphorylating mTORC1, ULK1, and autophagy-related proteins in the PIK3C3 complex[9]. Consequently, monitoring mTORC1 signaling flux is essential for investigating autophagy in trophoblast cells.

Hepatitis E virus (HEV) is a non-enveloped, icosahedral virus with a single-stranded positive-sense RNA genome. The viral particles measure approximately 32-34 nm in diameter, containing a 7.2 kb genome that features a 5’ methylguanosine cap and 3’ poly(A) tail[10]. The genome encodes four open reading frames (ORFs) - ORF1, ORF2, ORF3 and ORF4, with ORF4 being expressed exclusively in genotype 1 HEV. ORF1 contains multiple functional domains including methyltransferase (MeT), papain-like cysteine protease (PCP), hypervariable region (HVR), RNA helicase (Hel) and RNA-dependent RNA polymerase (RdRp)[11], primarily encoding non-structural proteins with well-characterized functions. ORF2 encodes the immunogenic capsid protein capable of inducing neutralizing antibodies. This protein is conserved among the four major pathogenic genotypes (HEV-1 to HEV-4), and the currently licensed HEV vaccine was developed using a modified recombinant antigen derived from HEV-1 ORF2 [12,13]. The ORF2-encoded capsid monomer consists of three distinct domains: S (shell), M (middle) and P (protruding). ORF3, which partially overlaps with ORF2, encodes a small multifunctional protein (∼12 kDa) involved in viral egress and quasi-envelope formation. This protein contains two hydrophobic domains (D1 and D2) and two proline-rich domains (P1 and P2), enabling the formation of membrane-associated oligomers through its proline-rich regions[14,15]. HEV infection generally manifests as self-limiting hepatic injury with subsequent complete clinical recovery and viral elimination, however, pregnant women exhibit markedly increased susceptibility to severe complications, including premature delivery, abortion and impaired fetal development[16].

Tophoblast cells represent the most functionally important cell type in the placenta[17], playing critical roles in pregnancy maintenance and embryonic development. Trophoblasts secrete various bioactive substances including steroid hormones, neuropeptide hormones, growth factors, and cytokines[18]. Moreover, the invasive migratory behavior of trophoblasts is essential for proper placental formation, embryonic development, and successful pregnancy progression[19]. Dysregulation of autophagic flux in trophoblasts significantly contributes to pregnancy disorders[20,21]. Tan et al. demonstrated that autophagy inhibition in trophoblasts markedly impairs their invasive capacity, with significantly suppressed autophagy observed in trophoblasts from spontaneous abortion cases[2]. Aiko et al. reported that ATG7 knockout in murine trophoblasts, leading to autophagy deficiency, resulted in placental hypoplasia[22]. Furthermore, Cao et al. proved that ATG16L1-mediated autophagy suppression can induce preterm birth[23].

The previous studies have demonstrated that HEV infection of placental trophoblast cells (JEG-3) leads to impaired autophagic flux, as evidenced by LC3B downregulation and p62 accumulation, resulting in defective autophagosome formation and ultimately abortion in ICR mice[24]. However, the precise molecular mechanisms underlying HEV-mediated autophagy suppression remain to be elucidated. Here, we successfully generated recombinant plasmids encoding HEV ORFs, using a combination of electron microscopy, Western blot, RT-PCR, and co-immunoprecipitation to systematically investigate the modulatory effects of HEV infection on host cell autophagy pathways. The study aims to identify viral proteins and critical functional domains responsible for autophagy inhibition, delineate the involved signaling mechanisms, and evaluate the therapeutic potential of targeting autophagy as a novel strategy for preventing and treating HEV infection during pregnancy in both humans and animals.

## Result

### Overexpression of HEV ORF2 and ORF3 in JEG-3 cells

The eukaryotic expression plasmid vectors used for transfection (Flag-ORF2_pcDNA3.1(+)-N-eGFP and Flag-ORF3_pcDNA3.1(+)-N-eGFP) carry the NeoR gene, conferring significant resistance to neomycin in successfully transfected cells. Therefore, G418 (Geneticin) can be used for cell selection to retain only the successfully transfected JEG-3 cells. The cell viability of JEG-3 cells under different G418 concentrations (0–1000 μg/mL) was determined by CCK-8 assay. The results showed that at a G418 concentration of 500 μ g/mL, untransfected JEG-3 cells (those without the resistance gene plasmid) were completely eliminated (cell viability < 30%). This concentration was thus determined as the optimal G418 screening concentration for post-transfection selection (S1 Fig).

Following drug selection, transfected JEG-3 cells were seeded onto coverslips for immunofluorescence analysis. As shown in Fig 1A, the overexpressed viral proteins exhibited distinct subcellular localization patterns due to their fused eGFP fluorescence signals. ORF2 primarily localized to the perinuclear region and nucleus. ORF3 was distributed throughout the cytoplasm, with significantly higher expression levels than ORF2. The difference in expression correlates with protein size, as larger proteins typically exhibit lower transfection efficiency compared to smaller ones.

**Fig 1.**
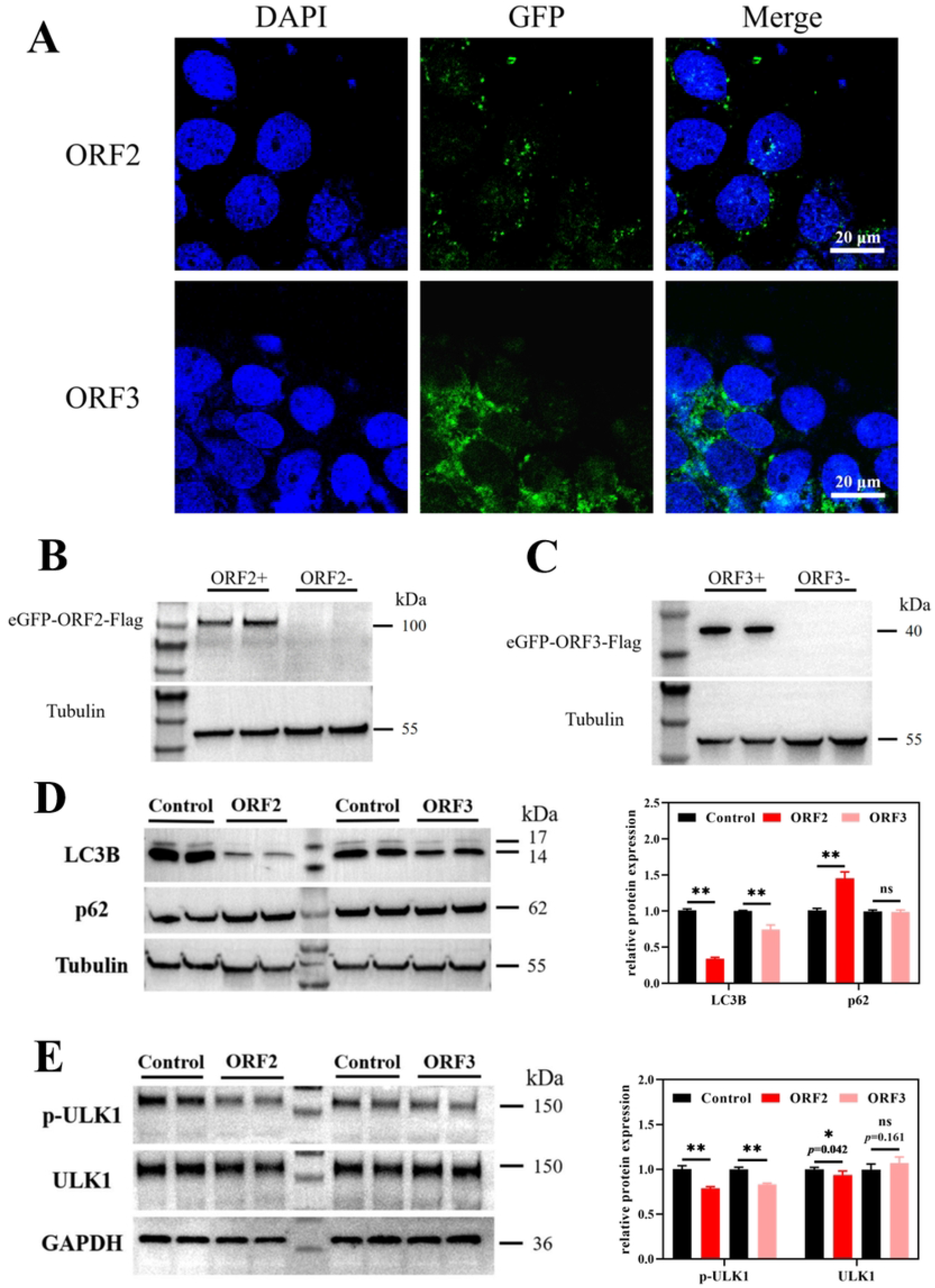
HEV ORF2 and ORF3 were successfully overexpressed in JEG-3 cells. Proteins and genes expression of LC3B and ULK1 in JEG-3 cells overexpressing HEV ORF2 and ORF3. (A)The JEG-3 cells transfected with ORF2 and ORF3 plasmids were observed under fluorescence microscope. The nucleus was stained with DAPI (blue), overexpressed ORF2 and 3 proteins were labeled with eGFP fluorescent protein signals (green). Scale bars represent 20 μm. (B) WB result of eGFP-ORF2-Flag and (C) eGFP-ORF3-Flag protein expression in JEG-3 cells (Flag antibody, 1:2000 dilution). eGFP protein is about 25kD and the size of Flag tag protein can be negligible. Therefore, the target protein size is about 100kD and 40kD respectively. (D) The expression of LC3 and p62 protein were significantly decreased in JEG-3 cells when HEV ORF2 protein was overexpressed, exhibiting typical autophagy inhibition. When HEV ORF3 protein was overexpressed, only LC3 protein expression decreased significantly. Bars indicate mean ± SEM, n= 6. **p < 0.01. (E) The expression of p-ULK1 protein were significantly decreased in JEG-3 cells when HEV ORF2 and ORF3 proteins were overexpressed. HEV ORF2 protein further significantly inhibited the expression of ULK1 protein. Bars indicate mean ± SEM, n = 6. *p < 0.05, **p < 0.01.

To confirm proteins overexpression, Western blot (WB) analysis was performed using the Flag tag fused to both proteins (Fig 1 B&C). In ORF2-transfected cells, a band of ∼100 kDa (eGFP: ∼25 kDa + ORF2: ∼75 kDa) was detected. In ORF3-transfected cells, a band of ∼40 kDa (eGFP: ∼25 kDa + ORF3: ∼15 kDa) was observed. These results align with the predicted molecular weights based on the plasmid-encoded sequences, confirming successful overexpression of HEV ORF2 and ORF3 in JEG-3 cells.

### HEV ORF2 plays a key role in autophagy inhibition in JEG-3 cells by suppressing ULK1 phosphorylation

The inhibitory effects of HEV proteins on autophagy in JEG-3 cells were investigated using cells overexpressing either HEV ORF2 or ORF3, with cells transfected with the Flag_pcDNA3.1(+)-N-eGFP plasmid serving as negative controls. As shown in Fig 1D, the expression levels of the autophagy marker LC3B were significantly reduced in JEG-3 cells overexpressing either ORF2 or ORF3 (*p* < 0.01). Additionally, ORF2 overexpression led to significant accumulation of p62 (*p* < 0.01), indicative of strong autophagy suppression. ORF3 overexpression also reduced LC3 levels but did not cause significant changes in p62. These results demonstrate that both HEV ORF2 and ORF3 inhibit autophagy in JEG-3 cells, with ORF2 exerting a more pronounced suppressive effect than ORF3. It suggests that HEV-induced autophagy inhibition in trophoblasts may be primarily mediated by ORF2.

The previous study demonstrated that HEV infection downregulates autophagic flux in chorionic trophoblast cells (JEG-3), suppressing all three stages of autophagy: initiation, progression and maturation. This suggests that HEV likely interferes with upstream autophagy-related signaling pathways through its viral proteins, leading to global inhibition of autophagic flux. To test this hypothesis, the expression of the autophagy initiation complex protein ULK1 and its phosphorylated form (p-ULK1) was examined in trophoblast cells overexpressing HEV ORF2 or ORF3. As shown in Fig 1E, both ORF2 and ORF3 significantly inhibited ULK1 phosphorylation (p-ULK1, *p* < 0.01). ORF2 additionally suppressed total ULK1 protein expression (*p* < 0.05). These findings indicate that HEV ORF2 and ORF3 block autophagy initiation by suppressing ULK1 phosphorylation, thereby disrupting subsequent autophagic processes. Notably, ORF2 exhibited stronger inhibitory effects on ULK1 than ORF3, consistent with its more pronounced suppression of LC3B observed earlier. Collectively, these results demonstrate that ORF2 likely serves as the primary mediator of HEV-induced autophagy suppression in JEG-3 cells.

### HEV ORF2 suppresses the upstream PI3K/Akt/mTOR pathway in JEG-3 cells

HEV significantly inhibits autophagy initiation and progression in JEG-3 cells. Since the PI3K/Akt/mTOR signaling pathway serves as a central negative regulator of autophagy, the study investigated whether HEV ORF2 targets this upstream regulatory axis. Based on previous findings demonstrating that ORF2 markedly reduces protein levels of the autophagy marker LC3 and the initiation complex ULK1, further examination was conducted on its effects on PI3K/Akt/mTOR signaling. As presented in Fig 2A&B, ORF2 significantly suppressed phosphorylation of PI3K and Akt (*p* < 0.01). Downstream mTOR and phospho-mTOR levels were consequently reduced (Fig 2C; *p* < 0.01), reflecting cascade inhibition of the entire pathway. Paradoxically, while PI3K/Akt/mTOR suppression typically activates autophagy, ORF2-mediated inhibition of this pathway coincides with overall autophagy repression in HEV-infected cells.

**Fig 2.**
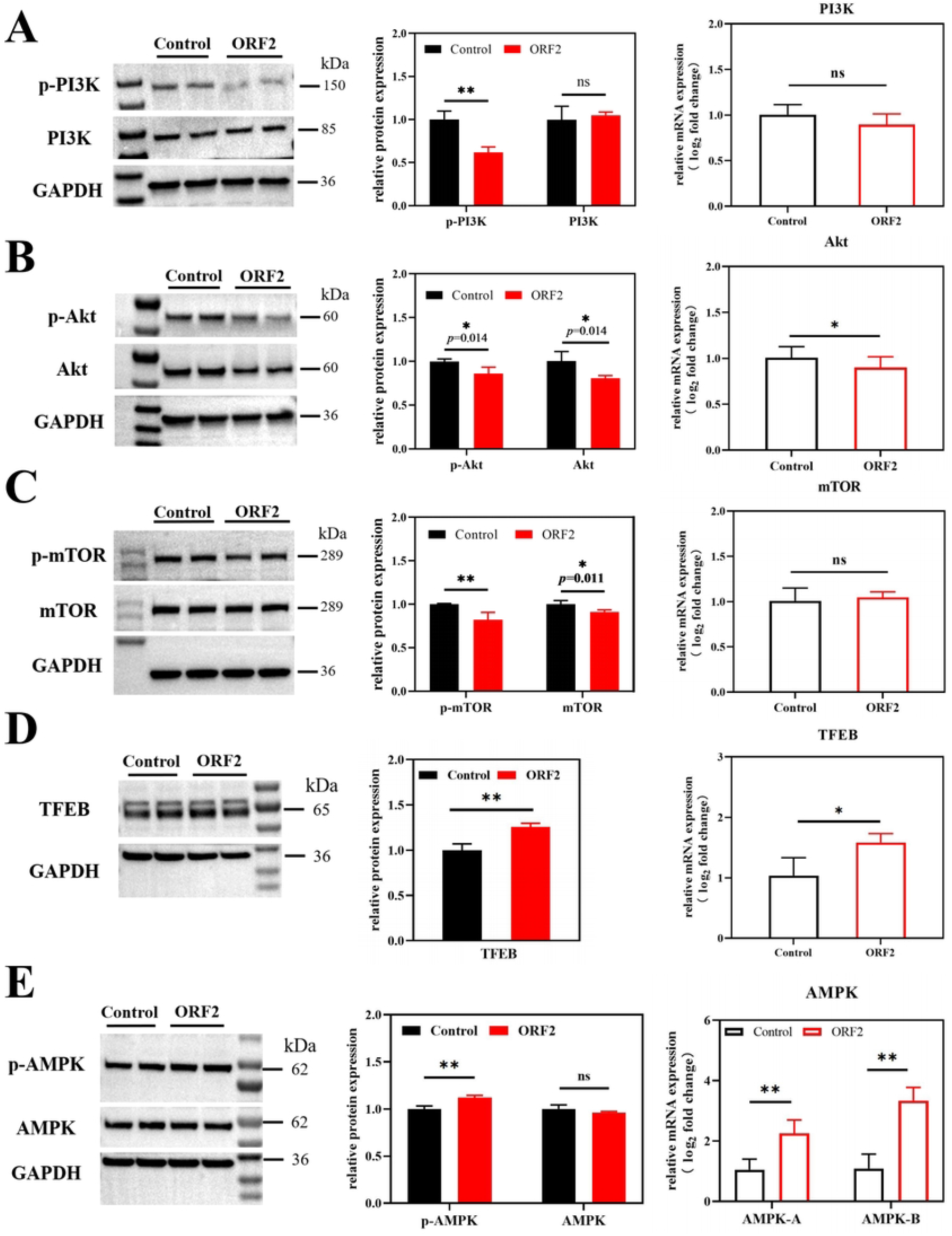
Proteins and genes expression of PI3K/Akt/mTOR, TFEB and AMPK in JEG-3 cells overexpressing HEV ORF2. (A) WB results and quantitative protein analysis of PI3K and p-PI3K. mRNA expression levels of PI3K. (B) WB results and quantitative protein analysis of Akt and p-Akt. mRNA expression levels of Akt. (C) WB results and quantitative protein analysis of mTOR and p-mTOR. mRNA expression levels of mTOR. (D) WB results, quantitative protein analysis and mRNA expression levels of TFEB. (E)WB results and quantitative protein analysis of AMPK and p-AMPK. mRNA expression levels of AMPK.

The interaction between the nuclear regulatory transcription factor EB (TFEB) and mTOR represents another core determinant of autophagy activation. Expression changes of TFEB, an mTOR-interacting protein, were therefore examined. As shown in Fig 2D, HEV ORF2 overexpression in JEG-3 cells significantly increased both transcriptional and protein levels of TFEB (*p* < 0.01). When considered alongside the observed inhibition of mTOR phosphorylation, these findings would typically indicate enhanced autophagic activity. However, this apparent pro-autophagic signature contrasts sharply with the strong suppression of LC3 protein in the autophagic flux, suggesting a paradoxical regulatory mechanism.

### HEV ORF2 activates AMPK to stimulate alternative autophagy pathways

AMPK, a central regulator of cellular energy metabolism, also activates autophagy through both direct and indirect mechanisms. Beyond the canonical mTOR-dependent pathway, AMPK regulates multiple alternative autophagy routes and positively modulates downstream autophagic processes. To investigate this compensatory mechanism, AMPK and phospho-AMPK expression were analyzed in JEG-3 cells overexpressing HEV ORF2. As shown in Fig 2E, ORF2 significantly upregulated both AMPK gene expression and p-AMPK protein levels (*p* < 0.01). This AMPK activation would typically indicate enhanced upstream autophagic signaling, yet paradoxically coincides with downstream autophagy suppression (evidenced by reduced LC3 levels).

### HEV ORF2 directly binds LC3 to suppress autophagy in JEG-3 cells

Although HEV ORF2 significantly suppresses the PI3K/Akt/mTOR signaling pathway and promotes AMPK phosphorylation - changes that would normally trigger autophagy activation - an unexpected suppression of autophagic activity was observed. These paradoxical findings suggest that the strong inhibition of LC3 protein and downregulation of ULK1 and its phosphorylated forms in ORF2-overexpressing JEG-3 cells may result from direct binding beguretween the viral ORF2 protein and host LC3.

Intermolecular interaction between HEV ORF2 and LC3 proteins was predicted using AlphaFold 3. As shown in Fig 3A, the ORF2 protein could theoretically bind directly to the LC3 protein, forming hydrogen bonds for stable linkage. In the structural prediction, valine at position 361 (V361) and glutamic acid at position 363 (E363) of the ORF2 protein may form hydrogen bonds with lysine at position 51 (K51), lysine at position 49 (K49), and arginine at position 70 (R70) of the LC3 protein. Additionally, asparagine at position 359 (N359), glycine at position 406 (G406), and arginine at position 542 (R542) of the ORF2 protein may form hydrogen bonds with leucine at position 53 (L53), histidine at position 27 (H27), and histidine at position 57 (H57) of the LC3 protein, respectively. To further verify the direct binding interaction between HEV ORF2 and LC3 proteins, a eukaryotic expression plasmid vector carrying red fluorescent signal (mcherry-LC3) was constructed and co-transfected with green fluorescent protein-tagged ORF2 and ORF3 plasmids (eGFP-ORF2/3) into JEG-3 cells. The distribution of red and green fluorescence was observed using fluorescence microscopy. Fig 3B shows that HEV ORF2 protein can co-localize with LC3 protein, with red and green fluorescence overlapping to form yellow fluorescent signals.

**Fig 3.**
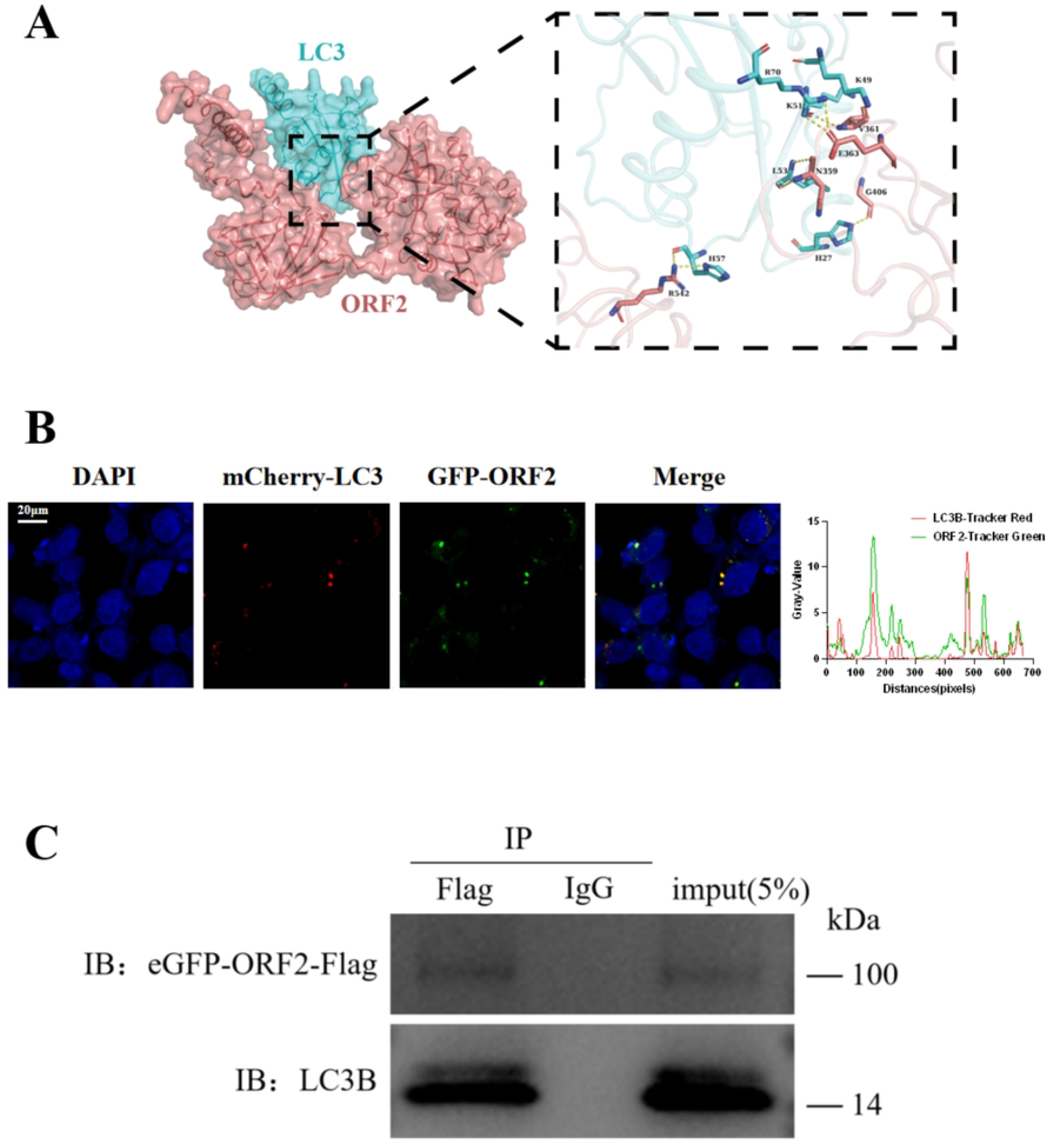
HEV ORF2 combined LC3 protein to inhibit autophagy of JEG-3 cells. (A) Prediction pattern graph of ORF2 and LC3 interaction structure using Alpha fold 3. LC3 (cyan) and ORF2 protein (pinkish lotus) structures are displayed in surface form. (B)The enlarged area is the interaction interface between LC3 and ORF2, where residues are shown as sticks and hydrogen bonds are shown as yellow dotted lines. (C) Immunoprecipitation (IP) was performed using anti-Flag antibody, followed by immunoblotting with anti-LC3B antibody. The ORF2 protein specifically co-precipitated with LC3B, while no interaction was detected in the IgG isotype control. Input lanes represent 5% of total lysates used for IP.

The interaction between HEV ORF2 protein and LC3 was validated using lysates from JEG-3 cells transfected with Flag-tagged ORF2 plasmid. The lysate supernatant was incubated with anti-Flag magnetic beads at 4°C overnight, using IgG antibody magnetic beads as an isotype control. After thorough washing, the immunoprecipitated complexes were eluted and detected by Western blot using anti-LC3 antibody. The results showed (Fig 3C) that a clear LC3 band could be detected in the ORF2 overexpression group, while no binding signal was observed in the IgG group. In conclusion, these results demonstrate that HEV ORF2 can inhibit trophoblast cell autophagy through direct binding with LC3 protein.

## Discussion

HEV ORF2 and ORF3 proteins were successfully overexpressed in JEG-3 cells. The results suggest that both HEV ORF2 and ORF3 proteins could significantly downregulate the expression of autophagy marker protein LC3B. Moreover, ORF2 caused a strong accumulation of p62, leading us to conclude that ORF2 is the key protein responsible for autophagy inhibition in JEG-3 cells. The previous research demonstrated that HEV inhibits all stages of autophagic flux (initiation, progression and maturation), suggesting that HEV’s mechanism of host cell autophagy inhibition targets upstream regulatory signaling pathways of autophagy. However, contrary to experimental expectations, HEV ORF2 significantly inhibited the activation of the core upstream autophagy regulatory pathway PI3K/Akt/mTOR while simultaneously upregulating both AMPK and TFEB pathways - theoretically this should, similar to other hepatitis viruses, activate downstream autophagy in cells. Ultimately, through AlphaFold 3, fluorescence co-localization experiments and Co-IP assays, it theoretically and experimentally demonstrated that HEV ORF2 can directly bind to autophagy marker protein LC3, thereby blocking the initiation and development of intracellular autophagy.

Interestingly, although ORF2 significantly inhibited PI3K/Akt/mTOR and activated the AMPK pathway, which should theoretically upregulate ULK1 expression, the experimental results is opposite. The regulation of autophagy by ULK1 is intricate. The ULK1 complex consists of activated ULK1, ATG13, ATG101, and FIP200, which activates the downstream Beclin-1 complex to initiate autophagy[25]. ULK1 can be degraded via the ubiquitin-proteasome pathway, and downregulation of p32 can significantly shorten ULK1’s half-life, reducing its protein levels[26]. Additionally, excess ULK1 may form small but dense autophagosome-like structures with Atg13, Atg14, Atg16 and other proteins, failing to properly activate autophagy and instead being degraded[27]. These findings suggest that the reduction in ULK1 expression may involve ubiquitin-proteasomal degradation.

HEV ORF2 encodes the viral capsid protein, which exhibits strong immunogenicity and is relatively conserved among the four major genotypes infecting humans, making it a primary target for vaccine development[28]. Currently, the commercially available HEV-1 ORF2-based vaccine Hecolin shows relatively weaker protection against HEV-3 and HEV-4, indicating structural differences in ORF2 proteins across genotypes[29]. Furthermore, phylogenetic clustering studies based on ORF2 sequences suggest that HEV-3ra differs significantly from HEV-3 and HEV-4 and should be classified separately[30]. These differences may explain why HEV-3ra, unlike other HEV genotypes, inhibits autophagy in JEG-3 cells instead of activating it as observed in hepatocytes. Recent studies have identified two forms of ORF2 in host cells: non-glycosylated ORF2 (ORF2c), which forms the viral capsid, and secreted ORF2 (ORF2s)[31]. ORF2s is abundant in cells and mediates virus-host interactions, contributing to immune evasion, viral replication support, and other functions. It is likely the primary form that binds host LC3 protein.

LC3B is cleaved by Atg4 protease, and its C-terminus covalently binds to phosphatidylethanolamine (PE), promoting membrane extension and autophagosome maturation, serving as a key indicator of autophagy activity. LC3B interacts with multiple autophagy receptor proteins, such as p62/SQSTM1, NBR1, and OPTN (Optineurin), through the LC3-interacting region (LIR)[32]. Current research shows that some pathogen proteins can directly and competitively bind to LC3B to inhibit autophagy. For instance, the nucleocapsid protein (NP) of Hantaviruses competes with viral glycoprotein (Gn) for LC3B binding, preventing autophagosome formation[33]. Influenza A virus protein PB1-F2 may physically interact with LC3B, inhibiting LC3B-mediated autophagosome formation and thereby impairing innate immunity[34]. Additionally, the nonstructural protein 1 of Respiratory syncytial virus (RSV) can disrupt normal cellular functions by binding to LC3B[35]. HEV ORF2 directly binds to LC3B in JEG-3 cells to inhibit autophagy, providing a novel drug target for HEV-induced placental damage and fetal miscarriage, as well as further evidence for the pathogenic mechanisms by which viral proteins directly bind LC3B.

In conclusion, the findings demonstrate that the binding of HEV ORF2 protein to LC3B serves as a key mechanistic target in host cell autophagy regulation. This discovery provides a theoretical foundation in autophagy research for elucidating the correlation between HEV infection and female reproductive system damage, as well as adverse pregnancy outcomes in pregnant animals/women.

## Materials and methods

### Plasmids

The eukaryotic expression plasmids used in this study, pcDNA3.1(+)-N-eGFP and pcDNA3.1(+)-N-6His, were synthesized and purchased from GenScript Biotech (Nanjing, China). Schematic diagrams of the two plasmid vectors are shown in S2 Fig, while the restriction enzyme sites and inserted protein sequences for each plasmid are detailed in S1 Table.

### Materials

The JEG-3 cell line was purchased from ATCC (Manassas, VA, USA). Reagents included: Minimum Essential Medium (MEM) (Gibco, Thermo Fisher Scientific), fetal bovine serum (FBS) (Gibco), phosphate-buffered saline (PBS) (Gibco), Lipofectamine 3000 (Invitrogen, Thermo Fisher Scientific), Opti-MEM (Gibco), G418 (Sigma-Aldrich, Cat# A1720), dimethyl sulfoxide (DMSO) (MCE, Cat# HY-Y0320), Trizol reagent (Thermo Fisher Scientific), RIPA lysis buffer (Solarbio, China), glycine (Beyotime Biotechnology), Tris-base powder (Beyotime Biotechnology), polyvinylidene difluoride (PVDF) membranes (Millipore), bovine serum albumin (BSA) (Beyotime Biotechnology), prestained protein ladder (Thermo Fisher Scientific), enhanced chemiluminescence (ECL) substrate (Millipore), antifade mounting medium with DAPI (Solarbio), 4% cell/tissue fixation solution (Solarbio), methanol, formaldehyde, xylene, absolute ethanol, isopropanol (Beijing Tongguang Fine Chemicals), chloroform (CAU Central Lab), and nuclease-free water (Solarbio). Antibodies used were: LC3B (ab192890, Abcam), p62 (Q13501, Abbkine Scientific), GAPDH (60004-1-1g, Proteintech), HRP-conjugated goat anti-mouse/rabbit IgG (Beyotime Biotechnology), Flag-tag (M20008, Abbkine Scientific), p-ULK1 (#6888, CST), ULK1 (#8054, CST), p-mTOR (#2971, CST), mTOR (#2983, CST), p-Akt (#4060, CST), Akt (#9272, CST), p-PI3K (#13857, CST), PI3K (#4257, CST), p-AMPK (TA3423, Abbkine Scientific), AMPK (T55326, Abbkine Scientific), TFEB (13372-1-AP, Proteintech), and α-Tubulin (M20005, Abbkine Scientific). Molecular biology reagents included reverse transcription premix kit (Accurate Biology) and SYBR Green master mix (Thermo Fisher Scientific).

### Plasmid transfection

Transfections were performed according to the manufacturer ‘ s protocol for Lipofectamine 3000 (Invitrogen). For standard transfection in 24-well plates, each well was transfected with 1 μ g plasmid DNA and 1.5 μ L Lipofectamine 3000 reagent, followed by 48 h of incubation. For co-transfection experiments involving LC3 with ORF2 or ORF3, each well received 0.5 μg mCherry-LC3 plasmid, 0.5 μg ORF2 or ORF3 plasmid, 3 μ L L3000 reagent, 2 μ L P3000 reagent, and 50 μ L Opti-MEM. After transfection, cells were seeded onto coverslips, and images were acquired immediately using a confocal microscope (A1HD25, Nikon).

### Cell selection

Since the transfected plasmids carried a kanamycin/neomycin resistance gene, transfected cells were selected using G418 (Geneticin, Sigma-Aldrich, Cat# A1720). To determine the minimum lethal concentration (MLC) of G418 for JEG-3 cells, a gradient of G418 concentrations (0, 50, 100, 200, 400, 500, 600, 800, 900, and 1000 μg/mL) was tested. JEG-3 cells were seeded in 96-well plates at a density of 5000 cells per well and allowed to adhere. The culture medium was then replaced with G418-containing medium, and cells were incubated at 37 ° C for 7-10 days. Fresh G418-supplemented medium was replenished every 3 days to maintain selection pressure. Once significant differences in cell viability were observed between groups, CCK-8 reagent (C0038, Beyotime Biotechnology) was added to quantify cell proliferation.

### RT-PCR

Trizol, provided by Invitrogen (Carlsbad, California, USA), was applied to extract RNA from JEG-3 cells. RNA was reverse-transcribed into cDNA and amplified with the kit mentioned in Materials, and the RT-PCR was performed using an ABI 7500 Real-Time PCR System (Thermo Fisher Scientific, USA). The primers used for autophagy-related genes in RT-PCR are listed in Supplementary Table 2.

### Western blot

Proteins were extracted from JEG-3 cells using RIPA lysis buffer (Solarbio, China) with 20 minutes of ice incubation. Protein concentrations were determined and normalized across experimental groups using the BCA assay kit (Thermo Fisher, USA). Prior to electrophoresis, samples were mixed with loading buffer (Cell Signaling Technology, USA) and denatured by boiling for 10 minutes. Electrophoretic separation was carried out on 4-20% Bis-Tris gradient gels followed by electroblotting onto PVDF membranes (Millipore, USA) at 200 mA constant current for 50-100 minutes, with transfer duration optimized according to target protein molecular weights. Membranes were blocked for 1 hour at room temperature with 5% non-fat dry milk (Mengniu, China) prepared in PBST solution. Primary antibody incubations were conducted overnight at 4 °C with 1:1000 dilutions. Following three PBST washes, membranes were probed with species-appropriate HRP-conjugated secondary antibodies (1 hour, room temperature). Protein bands were visualized using enhanced chemiluminescence substrate (Millipore, USA) and imaged with a Tanon 5200 detection system (China).

### Co-Immunoprecipitation

The Co-IP assay was performed using an immunoprecipitation kit (Abmart, #A10022) following the manufacturer’s instructions. The target protein was immunoprecipitated with an anti-Flag antibody (M20008, Abmart), with normal IgG antibody (Beyotime Biotechnology) serving as an isotype control. After overnight antibody incubation, 20 μ L of Protein A/G beads were added and incubated at 4°C with gentle rotation overnight. The samples were then centrifuged at 12,000 × g for 1 min, and the pellets were retained. The beads were washed three times with wash buffer. The immunoprecipitated complexes were resuspended in SDS loading buffer and denatured by boiling for 5 min. After centrifugation, the supernatant was collected for subsequent Western blot analysis.

## Acknowledgements

This work was supported by the National Key Laboratory of Intelligent Tracking and Forecasting for Infectious Diseases under Grant [Project No. 2024NITFID].

## Conflict of Interest

The authors declare no competing financial interest.

## Figure legend

TOC: The schematic diagram of HEV ORF2 inhibiting autophagy flux.

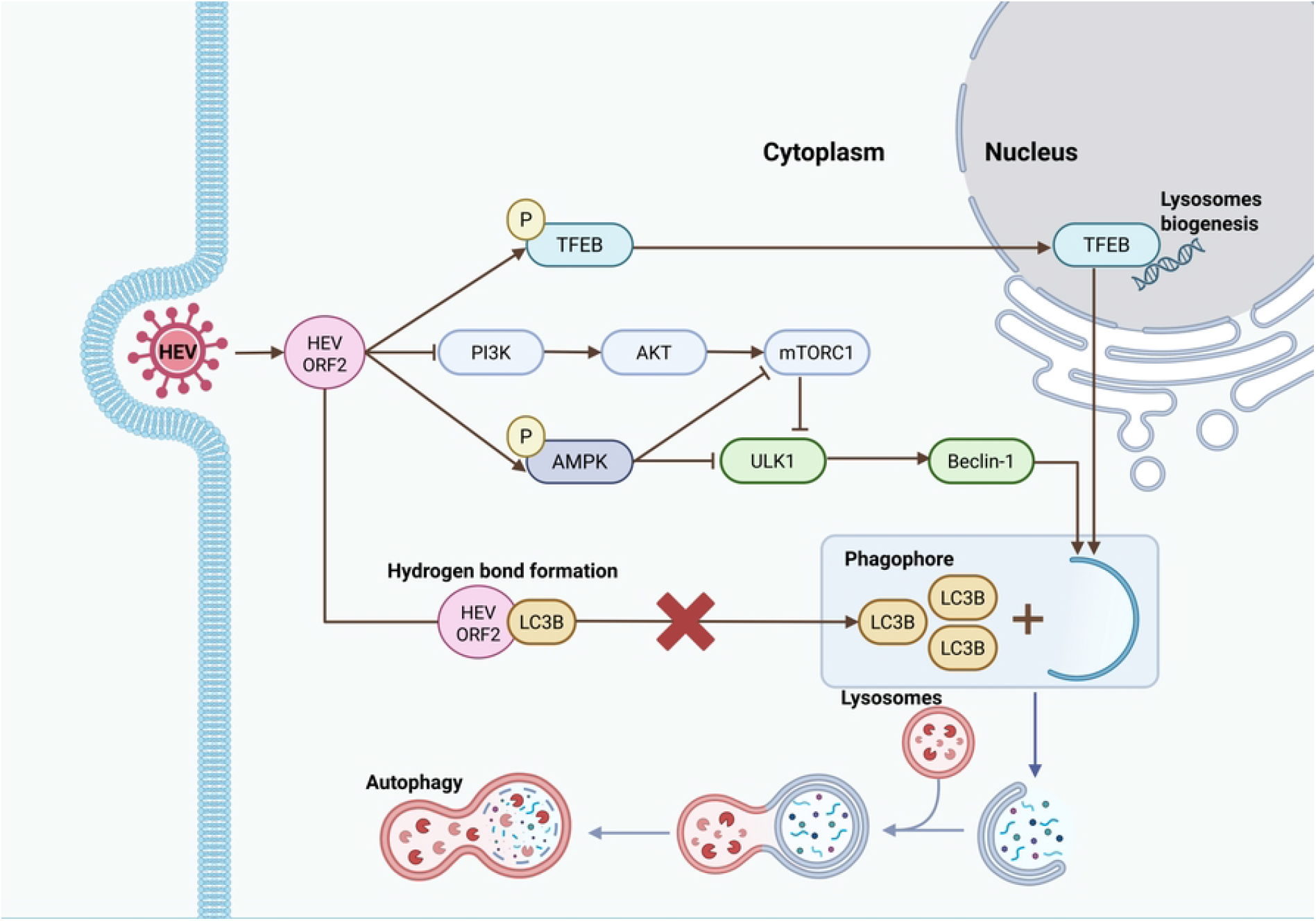

## Notes

### Competing Interest Statement

The authors have declared no competing interest.

## References

1. Feng Y, Chen Y, Wu X, Chen J, Zhou Q, Liu B, et al. Interplay of energy metabolism and autophagy. Autophagy. 2024 Jan;20(1):4–14.

2. Tan HX, Yang SL, Li MQ, Wang HY. Autophagy suppression of trophoblast cells induces pregnancy loss by activating decidual NK cytotoxicity and inhibiting trophoblast invasion. Cell Commun Signal. 2020 May 12;18(1):73.

3. Xiang H, Zhang J, Lin C, Zhang L, Liu B, Ouyang L. Targeting autophagy-related protein kinases for potential therapeutic purpose. Acta Pharmaceutica Sinica B. 2020 Apr 1;10(4):569–81.

4. Li MY, Shen HH, Cao XY, Gao XX, Xu FY, Ha SY, et al. Targeting a mTOR/autophagy axis: a double-edged sword of rapamycin in spontaneous miscarriage. Biomed Pharmacother. 2024 Aug;177:116976.

5. Lee S, Shin J, Kim JS, Shin J, Lee SK, Park HW. Targeting TBK1 Attenuates LPS-Induced NLRP3 Inflammasome Activation by Regulating of mTORC1 Pathways in Trophoblasts. Front Immunol. 2021;12:743700.

6. Wang J, Zheng F, Wang D, Yang Q. Regulation of ULK1 by WTAP/IGF2BP3 axis enhances mitophagy and progression in epithelial ovarian cancer. Cell Death Dis. 2024 Jan 29;15(1):97.

7. Cui Z, Napolitano G, de Araujo MEG, Esposito A, Monfregola J, Huber LA, et al. Structure of the lysosomal mTORC1-TFEB-Rag-Ragulator megacomplex. Nature. 2023 Feb;614(7948):572–9.

8. Zheng W, Zhang Y, Xu P, Wang Z, Shao X, Chen C, et al. TFEB safeguards trophoblast syncytialization in humans and mice. Proc Natl Acad Sci U S A. 2024 Jul 9;121(28):e2404062121.

9. Nwadike C, Williamson LE, Gallagher LE, Guan JL, Chan EYW. AMPK Inhibits ULK1-Dependent Autophagosome Formation and Lysosomal Acidification via Distinct Mechanisms. Mol Cell Biol. 2018 May 15;38(10):e00023–18.

10. LeDesma R, Nimgaonkar I, Ploss A. Hepatitis E Virus Replication. Viruses. 2019 Aug 6;11(8):719.

11. Muñoz-Chimeno M, Cenalmor A, Garcia-Lugo MA, Hernandez M, Rodriguez-Lazaro D, Avellon A. Proline-Rich Hypervariable Region of Hepatitis E Virus: Arranging the Disorder. Microorganisms. 2020 Sep 15;8(9):1417.

12. Zhou Z, Xie Y, Wu C, Nan Y. The Hepatitis E Virus Open Reading Frame 2 Protein: Beyond Viral Capsid. Front Microbiol. 2021;12:739124.

13. Peron JM, Larrue H, Izopet J, Buti M. The pressing need for a global HEV vaccine. J Hepatol. 2023 Sep;79(3):876–80.

14. Gouttenoire J, Pollán A, Abrami L, Oechslin N, Mauron J, Matter M, et al. Palmitoylation mediates membrane association of hepatitis E virus ORF3 protein and is required for infectious particle secretion. PLoS Pathog. 2018 Dec;14(12):e1007471.

15. Ding Q, Heller B, Capuccino JMV, Song B, Nimgaonkar I, Hrebikova G, et al. Hepatitis E virus ORF3 is a functional ion channel required for release of infectious particles. Proc Natl Acad Sci U S A. 2017 Jan 31;114(5):1147–52.

16. van Tong H, Hoan NX, Wang B, Wedemeyer H, Bock CT, Velavan TP. Hepatitis E Virus Mutations: Functional and Clinical Relevance. EBioMedicine. 2016 Sep;11:31–42.

17. Maltepe E, Fisher SJ. Placenta: the forgotten organ. Annu Rev Cell Dev Biol. 2015;31:523–52.

18. Turco MY, Gardner L, Kay RG, Hamilton RS, Prater M, Hollinshead MS, et al. Trophoblast organoids as a model for maternal-fetal interactions during human placentation. Nature. 2018 Dec;564(7735):263–7.

19. Abbas Y, Turco MY, Burton GJ, Moffett A. Investigation of human trophoblast invasion in vitro. Hum Reprod Update. 2020 Jun 18;26(4):501–13.

20. Yang HL, Lai ZZ, Shi JW, Zhou WJ, Mei J, Ye JF, et al. A defective lysophosphatidic acid-autophagy axis increases miscarriage risk by restricting decidual macrophage residence. Autophagy. 2022 Oct;18(10):2459–80.

21. Liu N, Shen H, Wang Z, Qin X, Li M, Zhang X. Autophagy Inhibition in Trophoblasts Induces Aberrant Shift in CXCR4+ Decidual NK Cell Phenotype Leading to Pregnancy Loss. J Clin Med. 2023 Dec 4;12(23):7491.

22. Sharma S. Autophagy-Based Diagnosis of Pregnancy Hypertension and Pre-Eclampsia. The American Journal of Pathology. 2018 Nov 1;188(11):2457–60.

23. Cao B, Macones C, Mysorekar IU. ATG16L1 governs placental infection risk and preterm birth in mice and women. JCI Insight. 2016 Dec 22;1(21):e86654.

24. Yang Y, Liu B, Tian J, Teng X, Liu T. Vital role of autophagy flux inhibition of placental trophoblast cells in pregnancy disorders induced by HEV infection. EMERGING MICROBES & INFECTIONS. 2023 Dec 8;12(2).

25. Lee DH, Park JS, Lee YS, Han J, Lee DK, Kwon SW, et al. SQSTM1/p62 activates NFE2L2/NRF2 via ULK1-mediated autophagic KEAP1 degradation and protects mouse liver from lipotoxicity. Autophagy. 2020 Nov;16(11):1949–73.

26. Jiao H, Su GQ, Dong W, Zhang L, Xie W, Yao LM, et al. Chaperone-like protein p32 regulates ULK1 stability and autophagy. Cell Death Differ. 2015 Nov;22(11):1812–23.

27. Banerjee C, Mehra D, Song D, Mancebo A, Park JM, Kim DH, et al. ULK1 forms distinct oligomeric states and nanoscopic structures during autophagy initiation. Sci Adv. 2023 Sep 29;9(39):eadh4094.

28. Aslan AT, Balaban HY. Hepatitis E virus: Epidemiology, diagnosis, clinical manifestations, and treatment. World J Gastroenterol. 2020 Oct 7;26(37):5543–60.

29. Sayed IM, Karam-Allah Ramadan H, Hafez MHR, Elkhawaga AA, El-Mokhtar MA. Hepatitis E virus (HEV) open reading frame 2: Role in pathogenesis and diagnosis in HEV infections. Rev Med Virol. 2022 Nov;32(6):e2401.

30. Khan N, Kakakhel S, Malik A, Nigar K, Akhtar S, Khan AA, et al. Genetic substructure and host-specific natural selection trend across vaccine-candidate ORF-2 capsid protein of hepatitis-E virus. J Viral Hepat. 2024 Sep;31(9):524–34.

31. Maas M, Neumann-Haefelin C. Deciphering the role of soluble ORF2 protein in virus-host interaction in HEV infection. Hepatology. 2023 Dec 1;78(6):1692–4.

32. Yao Y, Li S, Zhu Y, Xu Y, Hao S, Guo S, et al. miR-204 suppresses porcine reproductive and respiratory syndrome virus (PRRSV) replication via inhibiting LC3B-mediated autophagy. Virol Sin. 2023 Oct;38(5):690–8.

33. Wang K, Ma H, Liu H, Ye W, Li Z, Cheng L, et al. The Glycoprotein and Nucleocapsid Protein of Hantaviruses Manipulate Autophagy Flux to Restrain Host Innate Immune Responses. Cell Reports. 2019 May 14;27(7):2075-2091.e5.

34. Wang R, Zhu Y, Ren C, Yang S, Tian S, Chen H, et al. Influenza A virus protein PB1-F2 impairs innate immunity by inducing mitophagy. Autophagy. 2021 Feb;17(2):496–511.

35. Cheng J, Wang Y, Yin L, Liang W, Zhang J, Ma C, et al. The nonstructural protein 1 of respiratory syncytial virus hijacks host mitophagy as a novel mitophagy receptor to evade the type I IFN response in HEp-2 cells. mBio. 2023 Nov;14(6):e01480–23.

